# Genome-wide association study reveals *KCNQ3* gene as a risk factor for fear-related behavior in dogs and zebrafish

**DOI:** 10.1101/2025.05.07.652122

**Authors:** Eniko Kubinyi, Sára Sándor, Kitti Tátrai, Zsombor Varga, Zoltán Kristóf Varga, József Topál, Máté Varga, Dávid Jónás, Borbála Turcsán

**Affiliations:** Department of Ethology, ELTE Eötvös Loránd University, Budapest, Hungary; MTA-ELTE Lendület “Momentum” Companion Animal Research Group, Budapest, Hungary; ELTE NAP Canine Brain Research Group, Budapest, Hungary; Department of Genetics, ELTE Eötvös Loránd University, Budapest, Hungary; Institute of Experimental Medicine, Budapest, Hungary; HUN-REN Research Centre for Natural Sciences, Budapest, Hungary; ELTE – HUN-REN NAP Comparative Ethology Research Group, Budapest, Hungary

## Abstract

Freezing behavior, characterised by attentive immobility as a reaction to a perceived threat, is widely studied in the context of fear, anxiety and stress. To uncover the genetic factors underlying this behavior, we conducted a genome-wide association study (GWAS) in kennel-housed beagle dogs. Our analysis identified a single-nucleotide polymorphism (SNP) in intron 5 of the *KCNQ3* gene on chromosome 13 associated with freezing behavior in response to unfamiliar environments and people. To validate this finding, we used a zebrafish model, where CRISPR/Cas9-induced kcnq3 deficiencies led to heightened fear and arousal in two behavioral tests. *KCNQ3* is implicated in several neurodevelopmental and psychiatric disorders, and our results highlight its evolutionarily conserved role in modulating fear responses. In dogs, an enriched environment can mitigate the adverse effects of *KCNQ3* deficiency by reducing threat perception, highlighting the role of gene-environment interactions in shaping behavioral responses.

**Teaser:** *KCNQ3* influences fear responses across species, with gene-environment interactions shaping freezing behavior in dogs.

## Introduction

Freezing is a defensive behavior characterized by attentive immobility and increased muscle tone, allowing animals to avoid detection by predators. Unlike active responses such as fight or flight, which are mediated by the sympathetic nervous system, freezing is a passive survival strategy involving both sympathetic and parasympathetic activity. This strategy can be sustained for extended periods when escape is not an option (*1*–*3*). While evolutionary adaptive, excessive or inappropriate freezing is linked to mental disorders in humans, often as a consequence of early-life adversities (*4, 5*). Understanding the genetic and environmental factors influencing freezing behavior can provide critical insights into the mechanisms underlying stress-related psychopathologies (*6, 7*).

Dogs offer a valuable model for studying fear-related behaviors due to their close evolutionary and ecological relationship with humans. Their behavioral and environmental similarities to humans make them particularly relevant for translational psychiatric research (*8, 9*). Previous studies estimate that the heritability of fearful behavior in dogs is around 0.2 (*10*). Additional evidence for genetic contributions to fear-related behaviors comes from historical research on selectively bred pointer dogs, where individuals from “nervous strains” exhibited increased freezing, heightened avoidance responses, and reduced exploratory behavior in novel environments. Rehabilitation through increased human contact was ineffective, whereas social facilitation and targeted desensitization gradually alleviated these behaviors (*11, 12*). Genome-wide association studies (GWAS) have identified several genes associated with fear-related behaviors in dogs (*13*–*16*).

However, most previous GWAS were conducted on pet dogs with diverse environmental backgrounds, which may have introduced confounding factors that obscure genetic effects (*14*). For instance, associations between the dopamine receptor D4 gene (*DRD4*) and activity-impulsivity were detected in police dogs raised under controlled conditions but not in pet dogs (*17*). In a more recent study, 44% of kennel-housed beagles exhibited defensive behaviors, primarily freezing and flight, in a standardized test. In contrast, pet dogs from the same genetic lineage, as well as the offspring of kennel-housed dogs raised in enriched environments, showed no freezing (*18*). These findings highlight the mitigating effect of environmental enrichment on fear responses, emphasizing the importance of studying genetically similar populations under standardized conditions.

To address these limitations, we aimed to identify genetic factors underlying fear-related responses in dogs while minimizing environmental variability. We analyzed the behavior of 50 kennel-housed beagles in a standardized test battery consisting of 14 situations. Based on their physical activity and responsiveness, we classified the dogs into four behavioral clusters. We then conducted a GWAS to identify genetic loci associated with freezing behavior, followed by sequencing and gene expression analysis in brain tissues from affected and non-affected dogs. Finally, to validate our findings, we generated zebrafish (*Danio rerio*) strains deficient in the candidate gene and assessed their fear-related behaviors in two behavioral tests.

## Results

### Behavioral analysis and GWAS in dogs

A hierarchical cluster analysis (Ward methods) based on the two behavioral variables (activity and responsiveness) grouped the dogs into four clusters (Fig. 1, Fig. S1). Cluster 1 involved dogs showing freezing behavior, Cluster 4 the active-responsive dogs, Clusters 1-2 the intermediate types. Notably, the behavior of dogs in Cluster 4, despite being raised in kennels, was similar to pet dogs living in enriched environments, measured in Turcsán et al. (2020), where pet dogs’ total activity median was 56.4% and responsiveness median was 1.8.

**Figure 1.**
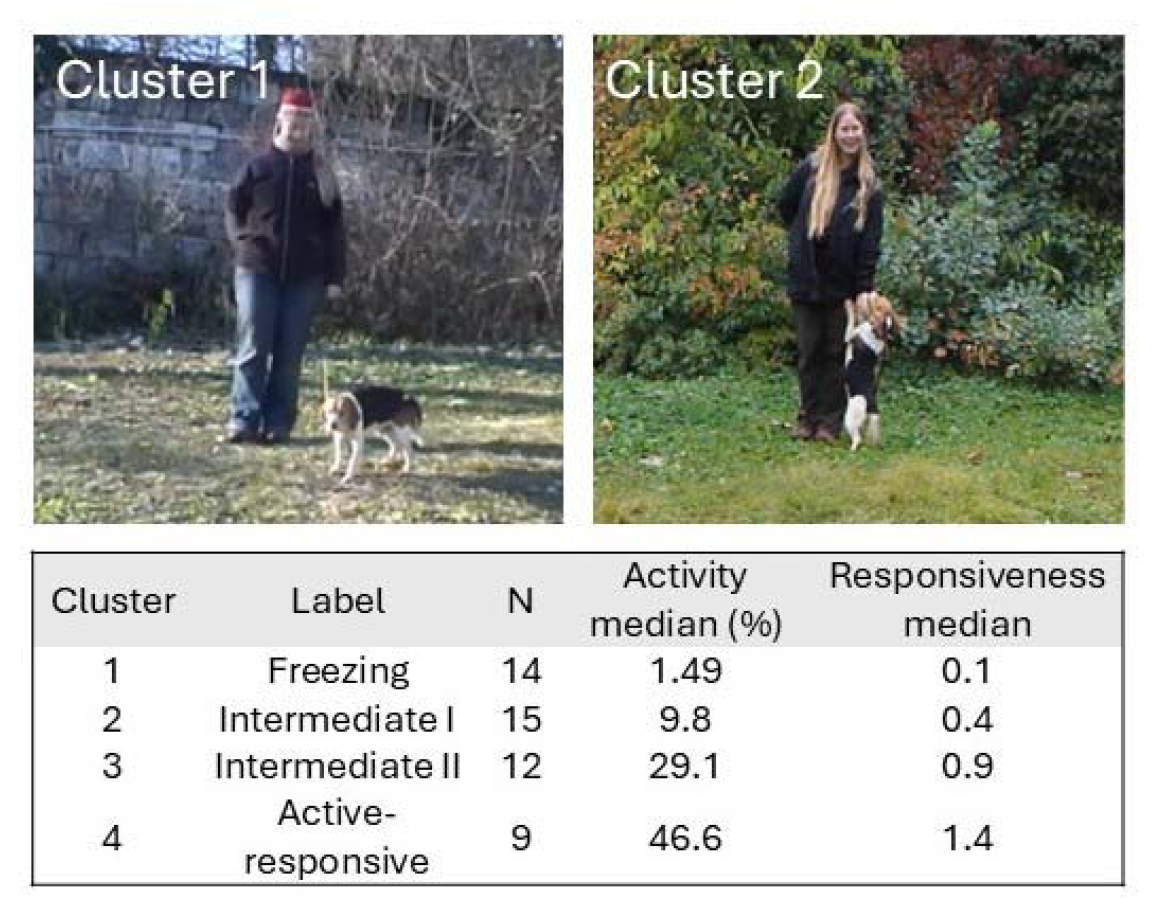
Phenotyping dogs. Two examples from Clusters 1 and 2 (top), and behavioral characteristics of the clusters. (The individual depicted is BT, one of the co-authors of the paper. She provided written consent for the publication.)

Comparing Cluster 1 vs 4, GWAS identified one significant SNP on chromosome 13 (Fig. S2) with a Bonferroni-corrected p-value of 6.35*10^−8^ (or -log□□(p) = 7.2), with sex as a covariate showing no significant effect. The identified QTL exhibited a p-value that was 5 million times lower compared to neighbouring SNPs. No other SNP on the SNP array was in complete LD with the detected QTL.

The significant SNP - a transversion-type mutation (C/G) can be found in the dbSNP database with the rs22253798 ID. It is located in the fifth intron of the *potassium voltage-gated channel subfamily Q member 3* (*KCNQ3*; Ensemble ID of the CanFam 3.1 reference genome: ENSCAFG00000001105, SNP position: 28974415). Although out of the three transcripts annotated for this gene (see Supplement Materials (SM)), only two transcripts encompass this section of the annotated gene body, the protein level annotation indicates that intron five separates two exons, which encode the ion transport domain of the protein, thus suggesting that the identified SNP marks a functionally relevant section of the gene.

*KCNQ3* is a protein-coding gene located on chromosome 13 with a total length of 291,434 base pairs (Fig. 2a). It consists of 22 exons in total and is transcribed from the reverse strand. Three transcripts are produced from it, namely *KCNQ3-201* (length: 751 amino acids (AA)), *KCNQ3-202* (length: 751 AA) and *KCNQ3-203* (length: 872 AA). Plassais et al. (2019) published 3703 mutations overlapping with this gene, most of which are intronic variants (Fig. 2b and 2c). The identified mutation is located in intron 5, which separates two exons that encode one of the two protein domains of the gene (ion transport domain), therefore overlapping with a part of the gene that is potentially functionally active.

**Figure 2.**
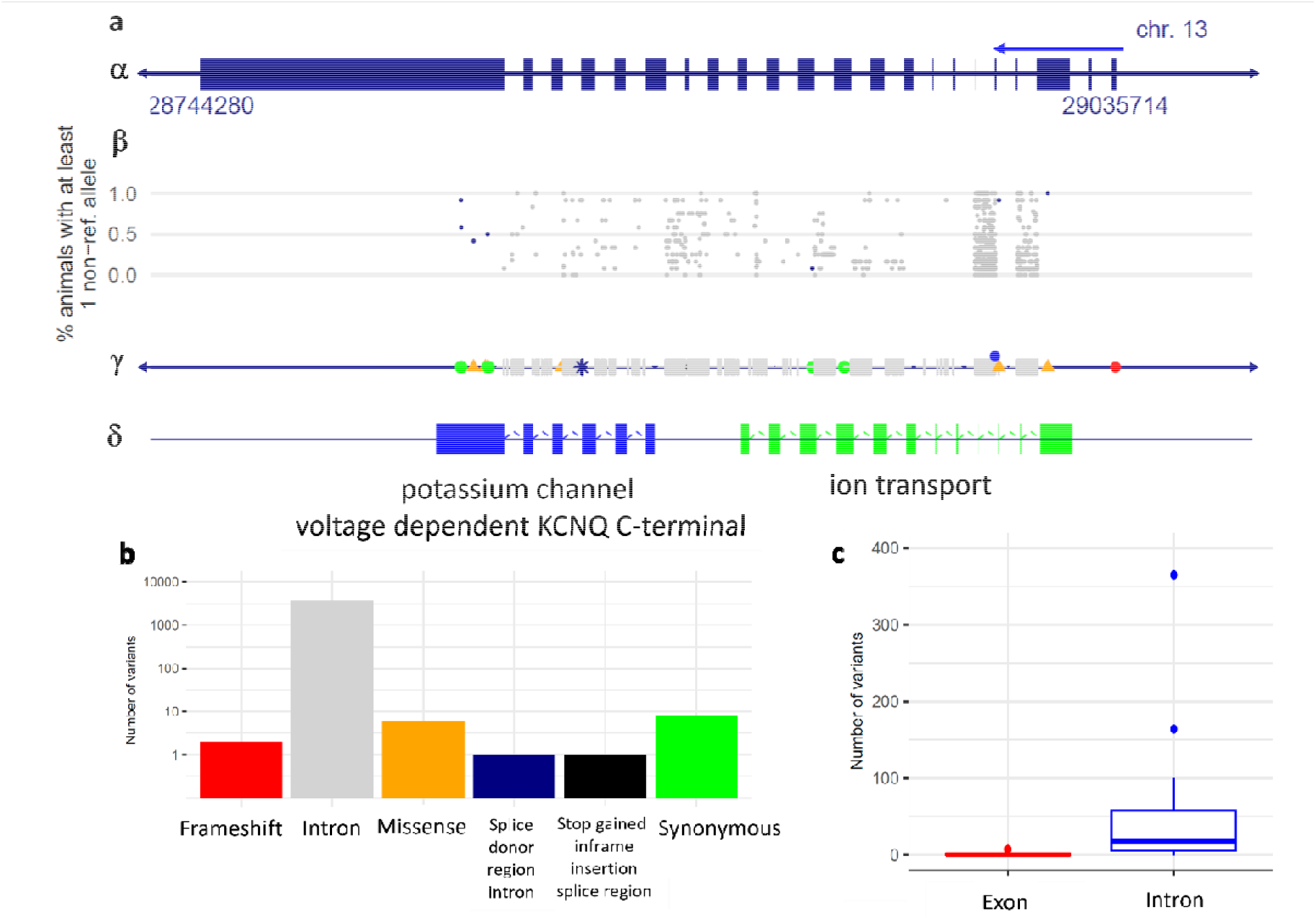
Genetic features of the *KCNQ3* gene. **(a)** *α*: Outline and genetic features of the *KCNQ3* gene. Notably, the genetic positions were lifted over to the Dog10K_Boxer_Tasha reference genome, due to the availability of protein domain regions only for this genome version. Exons are marked as larger blocks while introns and intergenic regions as narrow lines. The transcription direction is indicated by the blue arrow. All introns were narrowed down to 200 bp in size to make the exons visible. *β*: Proportion of scent hounds, including beagles, carrying at least one non-reference allele in genetic variations within *KCNQ3*. The scent hound breed group was defined based on Parker et al. (2017). Scent hounds with SNP data from Plassais et al. (2019) were selected for this analysis. *γ*: Distribution of the different known genetic variations across the *KCNQ3* gene, was annotated with Ensembl’s Variant Effect Predictor. Annotation categories and color coding are the same as in Fig. 1b. The significant mutation detected in this study is marked with a blue dot above intron 5. *δ*: Protein domain regions of *KCNQ3* predicted and published by the UniProt database. **(b)** Frequency distribution of different annotation categories. **(c)** Boxplots showing the distribution of genetic variations in exons and introns. Note that the longest intron contained 2703 variants, which is not visible on the boxplot.

### KCNQ3 expression and sequence analyses in dogs

Since the GWAS analysis implicated *KCNQ3* in the background of the observed phenotype, we investigated its expression and potential coding sequence (CDS) mutations by: (i) comparing *KCNQ3* expression in the frontal cortex between freezing and non-freezing dogs, (ii) sequencing frontal cortex RNA (cDNA), (iii) analyzing brainstem total RNA sequencing datasets, and (iv) sequencing a main region of interest within the *KCNQ3* gene using genomic DNA.

We found that the (i) expression level of *KCNQ3* in frontal cortex samples did not differ between the dogs (Fig. S3, Table S1). (ii) The RNA Sanger sequencing, in a subset of the frontal cortex samples of the “freezing” dog (Cluster 1 in Table S1) showed genomic mosaicism, and some cells carried an allele with a 33 amino acid long in-frame deletion in exon 1, L58_L90del, suggesting brain mosaicism. We identified one additional missense and three same-sense mutations (Table S2). Based on the prediction of the PROVEAN web server (*21*), the missense mutation is a neutral mutation; therefore, it is unlikely to affect the function of the protein. Neither (iii) brain stem total RNA sequence data (Table S3) nor (iv) DNA Sanger sequence data were different across dogs (Table S3, Fig. S4).

### *Validating the* KCNQ3*’s role in stress regulation in a fish model*

To explore a potential functional connection between *KCNQ3* and behavior, we extended our research to zebrafish (*Danio rerio*), comparing fear-related escape and anxiety-related avoidance responses in *kcnq3* knockout and wild-type individuals. The dog and zebrafish *KCNQ3* orthologs are 63% identical and 71% similar (Fig. S5). We sought to investigate whether the absence of *kcnq3* in “crispant” animals exerts changes in affective behavior.

We found significant effects of the time point (F=68.82, p<0.001), the genotype (F=19.914, p<0.001), and their interaction (F=25.456, p<0.001) on the startle response to the tactile stimuli, indicating differences in both the magnitude and the dynamics of escape responses (Fig. 3a; details of the statistical analysis are in SM). We also calculated cumulative measures for the 10 seconds before and after the first tactile stimulus to assess general changes in locomotion. We found that while the wild-type animals did not increase their activity apart from prompt responses to each stimulus (t=1.148, p=0.255), the *kcnq3*-impaired group exhibited generally heightened activity (t=3.937, p<0.001), a potential marker of an enhanced state of arousal and fear (Fig. 3b). In the SPM test, the knockout group showed an enhanced anxiety score, calculated from the frequency and latency to enter and the time spent in the shallow arm of the platform (t=2.663, p=0.01) (Fig. 3c). We did not find locomotor differences in the SPM (t=0.884, p=0.380), indicating that the measured changes in the shallow arm activity are not the consequence of altered general movement levels but are specific to anxiety-like behavior (Fig. 3d).

**Figure 3.**
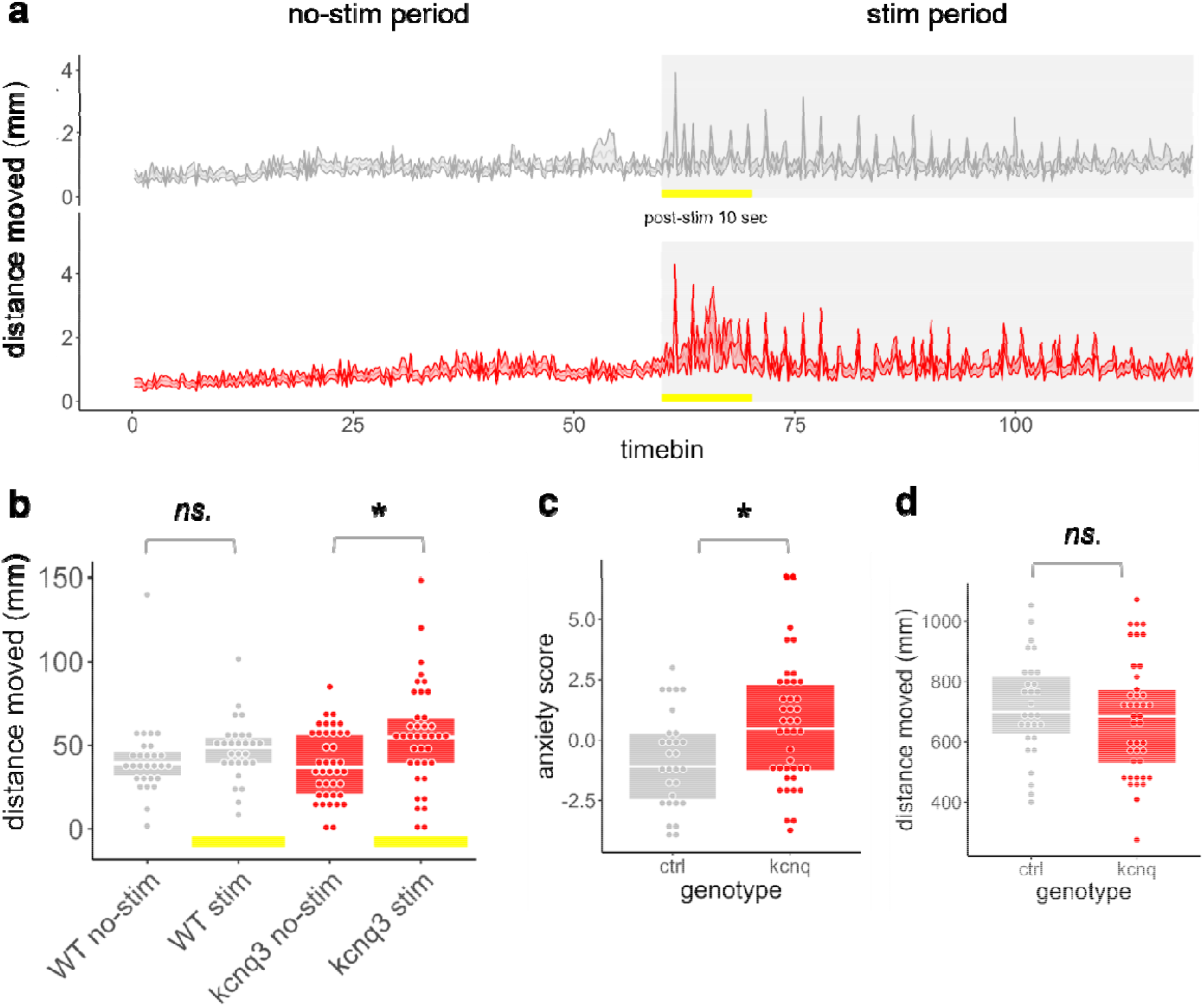
Behavioral response measurements in the Tactile Startle (TS) and Swimming Plus-Maze (SPM) tests. **(a)** Distance moved in the TS as a function of time and tactile stimuli. Grey and white backgrounds indicate periods with and without tactile stimuli, respectively. Yellow bars mark the 10-second periods used to quantify general locomotion changes in response to the stimulus. **(b)** Summarization of movement 10 seconds before (no-stim) and after (stim) the start stimulus onset. **(c)** Anxiety scores calculated from shallow arm activity -(scaled frequency + scaled time - scaled latency) in the SPM test, reflecting avoidance of shallow areas, an indicator of anxiety-like behavior in zebrafish. *indicates significant difference **(d)** Distance moved during the SPM test.

## Discussion

Our study identifies *KCNQ3* as a gene associated with freezing behavior in kennel-housed beagle dogs and enhanced arousal in zebrafish, suggesting a conserved role in regulating fear responses across vertebrate species. *KCNQ3* encodes a subunit of voltage-gated potassium channels, essential for neuronal excitability and the balance between excitation and inhibition. Dysregulation of these channels can contribute to hyperexcitability, a hallmark of several neurological and psychiatric conditions. Our findings align with previous research linking *KCNQ3* to brain excitability disorders, including seizures, intellectual disabilities, and epileptic encephalopathies (*22, 23*). Recent work in mice further supports *KCNQ3*’s role in anxiety when it was revealed that threat-induced anxiety can affect the expression of potassium channel genes (most prominently *kcnq3*) in the ventral bed nucleus of the stria terminalis (vBNST) (*24*). Therefore, misregulation of *KCNQ3* in the vBNST could result in excess anxiety and freezing.

Beyond genetics, our findings emphasize the importance of considering gene-environment interactions in the study of fear and anxiety. The absence of freezing behavior in family dogs from the same gene pool as the kennel-housed dogs (*18*) suggests that environmental enrichment, characterized by increased stimulation and social interactions, can buffer against genetic predispositions to maladaptive fear responses. This aligns with research suggesting that enriched environments promote neural plasticity and reduce stress-induced hyperexcitability, mitigating fear-related behaviors in dogs (*11*) and other animals (e.g., (*25*)).

However, our study has limitations. The small sample size increased the likelihood of false negatives in GWAS, while recombination hotspots near the candidate loci may have hindered the detection of true associations. Additionally, the lack of significant findings in our gene expression analysis is likely due to the limited number of available brain samples or the possibility that our methods were not optimized to capture fine-scale, transient changes in *KCNQ3* expression that may occur in response to acute threats in specific brain regions. Nevertheless, gene expression data from humans and mice indicate that *KCNQ3* is broadly expressed across the brain (*26, 27*).

Overall, our study provides evidence that a functional mutation in the *KCNQ3* gene influences fear and anxiety behaviors across vertebrates, underscoring the evolutionary conservation of *KCNQ3* in stress-response pathways. The lack of freezing behavior in family-adopted dogs representing the same gene pool but living in an enriched environment (*18*) highlights the interplay between genetic predispositions and environmental influences. The findings have translational implications for both human psychiatric research and practical applications for improving the welfare of kennelled dogs, where pathological fear remains a critical concern (*28*). Future research should prioritize validating *KCNQ3*’s role in stress-response mechanisms using larger sample sizes and broader gene expression analyses across diverse brain regions. This could facilitate the development of therapeutic interventions targeting KCNQ3 to address maladaptive fear and anxiety responses, benefiting both clinical and veterinary fields.

## Materials and Methods

### Phenotyping dogs

Fifty kennel-housed beagle dogs (74% males, mean age ± SD: 2.71 ± 1.42 years) were tested in a behavioral test battery, while on a 1.5–2 m leash, in a grassy area, 30 m away from the dogs’ kennels with two unfamiliar, friendly female experimenters, one playing the role of the owner. The test battery consisted of 14 subtests designed to assess dogs’ spontaneous activity, social interactions, problem-solving abilities, and responses to novel or potentially threatening stimuli. The test began with spontaneous exploration, followed by a greeting interaction where the experimenter attempted to pet the dog. Dogs’ attention and frustration were assessed using a pendulum test in which a sausage was swung just out of reach. Separation tolerance was evaluated across four phases, alternating between brief isolation and reunion with either the experimenter or the ‘owner”. Playfulness was measured through ball play, while problem-solving was tested by allowing dogs to retrieve food from a small cage. Resource guarding tendencies were examined by offering a large bone and then attempting to take it away. Defensive reactions were assessed using a threatening approach, and startle responses were measured by suddenly opening an umbrella near the dog. Compliance and restraint tolerance were tested by attempting to lay the dog on its side. Decision-making and susceptibility to human influence were evaluated in a food choice task where the experimenter demonstrated a preference for a smaller food portion. Attachment and seeking behavior were assessed in a hiding test, in which the ‘owner’ temporarily left the dog with the experimenter. Finally, spontaneous activity was measured again at the end of the test battery to compare activity levels before and after exposure to the various tasks (Fig. 1A, Fig. S6, Table S4, Video S1 or https://youtu.be/i2zJ0C-5k8w, Video S2 or https://youtu.be/shK3Nk4MAPU). All tests were video recorded. Based on the videos, two variables were obtained: activity and responsiveness scores. Activity is the percent of time dogs engaged in physical activity during the tests. Responsiveness score is calculated as the mean of 50 behavioral variables scored on a 0–3 scale in each subtest, where 0: no reaction, 3: intensive (positive) reaction (*18*). Freezing was characterized by low activity and low responsiveness, while active-responsive (sociable) dogs were characterized by high activity and high responsiveness. For more details, see Supplemental Materials (SM).

### Genotyping dogs

DNA was extracted from buccal swabs (*17*). Genotyping was performed using the Illumina 170K Canine HD SNP array (Illumina, San Diego, CA, USA), which included 169,136 SNPs. After SNP quality control, 82,409 SNPs were retained for the GWAS (see SM). The genomic inflation factor (λ = 1.01) and the Q-Q plot indicated no significant population stratification or systematic technical bias in the analysis (Fig. S7). We carried out two GWAS analyses. First, we compared only the two extreme phenotypes, Cluster 1 vs 4. Next, we compared Cluster 1+2 vs Cluster 3+4.

#### *KCNQ3* expression analysis

Brain samples (frontal cortex) from five dogs with known behavioral phenotypes (one freezing, one intermediate 1, one intermediate 2, and two active-responsive, Table S3) were used for total RNA isolation. The protocols for the RNA isolation, cDNA synthesis, qRT-PCR, and the primers (Table S5) are presented in the SM.

#### RNA Sanger sequencing

We used the same samples as in the previous study (Table S3). Standard qRT-PCR and Sanger sequencing were conducted on *KCNQ3* CDS clones generated from overlapping PCR fragments, cloned into *E. coli*, and verified using standard SP6 primers. Further details and primers (Table S6) are shown in the SM.

#### Total RNA sequencing

Total RNA sequence data of brain stem samples were available for four dogs (1 “freezing”, 1 “freezing-intermediate”, 1 “intermediate”, and 1 “active-responsive”, Table S2), originally sequenced for (*29*)17. Sequencing and analysis of the *KCNQ3* gene, including alignment to the CanFam v3.1 reference genome were performed as described in (*29*), with raw data accessible under Bioproject ID: PRJNA939639. We also calculated CPM (counts of reads per gene per million sequenced reads).

#### DNA Sanger sequencing

Genomic DNA was isolated from buccal swabs of nine dogs (five “freezing”, two “intermediate”, two “active-responsive”, Table S3 and Table S7), using a TNES buffer and ethanol precipitation method, followed by purification and resuspension in TE buffer, while genotyping was performed by amplifying a 232 bp portion of *KCNQ3* exon 1 using PCR and sequencing the fragments via Sanger sequencing. For more details, see SM. We found that the samples did not differ from the reference genome (ROS_Cfam_1.0).

### Phenotyping zebrafish

Zebrafish were maintained in the animal facility of ELTE Eötvös Loránd University according to standard protocols (*30*). Ten embryos (two days post fertilisation) and five cohorts of fish (30 days post fertilisation) were used in this study. To create zebrafish deficient (“crispant”) in *KCNQ3* we used recently developed protocols that can create efficient gene knock-down phenotypes in F0 animals using a CRISPR/Cas9-based methodology (*31, 32*). For details, see SM, Table S8.

Wild type and *kcnq3* “crispant” fish were tested simultaneously at 30 days post-fertilization in two tests in succession. We used five cohorts of zebrafish for the experiments. In the Tactile Startle (TS) paradigm, zebrafish were exposed to repeated tactile stimuli in a controlled environment, and their locomotor response (e.g., startle or escape) was analyzed to infer fear and arousal states. In the Swimming Plus-Maze (SPM) test, zebrafish were placed in a plus-shaped maze with shallow and deep arms, and their preference for shallow arms (indicative of anxiety-like behavior) was quantified using an anxiety score derived from latency, frequency, and time spent in the shallow arms (*33*). General locomotion was also measured in both tests. For further details, see SM.

Genomic DNA was extracted from 10 embryos at two days post fertilization, and after PCR amplification, Sanger sequencing was used to confirm the efficiency of targeting, which suggested the presence of multiple indels upon the injection of the Cas9 RNPs (Fig. S8). For target sequences and primers, see SM, Table S9.

## Acknowledgments

We would like to acknowledge the generous help of Elaine A. Ostrander and her laboratory for their contribution to the genotyping of dogs. Eszter Petró helped with the behavioral testing, and Zsófia Dobó, Tímea Vándor, and Henrietta Bolló helped with the behavioral coding of the dogs. DNA was extracted by the laboratory of the Department of Medical Chemistry, Molecular Biology and Pathobiochemistry, Semmelweis University. During the preparation of this work, the authors used Grammarly and ChatGPT for grammar correction. After using this tool/service, the authors reviewed and edited the content as needed and take full responsibility for the content of the publication.

## Funding

Hungarian Academy of Sciences MTA-ELTE ‘Lendület/Momentum’ Companion Animal

Research Group grant PH1404/21

National Brain Programme 3.0 grant NAP2022-I-3/2022

Hungarian Ethology Society

KIFÜ (the Hungarian Governmental Informatics Development Agency) granted access to high-performance computing resources

Hungarian Academy of Sciences (HAS) János Bolyai Fellow grant BO/00555/22/8

## Author contributions

Conceptualization: EK, JT, BT, MV

Methodology: EK, TB, JD, MV, SS

Investigation: EK, TB, JD, ZV, ZsV

Visualization: EK, BT, JD, ZV

Supervision: EK, MV

Writing—original draft: EK, BT, JD, MV, ZV

Writing—review & editing: EK, BT, JD, MV, ZV, SS, ZsV

## Data and materials availability

All data are available in the main text or the supplementary materials.

## Supplementary Materials

