## Supplementary Materials for "Genome-wide association study reveals *KCNQ3* gene as a risk factor for fear-related behavior in dogs and zebrafish"

Eniko Kubinyi *et al.*

**This PDF file includes:**

Supplementary Text

Figs. S1 to S8

Tables S1 to S9

Movies S1 to S2

Data S1

**Other Supplementary Materials for this manuscript include the following:**

Movies S1 to S#

Audio S1 to S#

Data S1 to S#

Supplementary Text

*Please note that in the Supplementary Text, we follow the Figure and Table numbering as they appeared in the Main text.*

### 1. Identifying risk loci associated with freezing behavior in dogs

#### Subjects

Subjects were kennel-housed beagle dogs (N = 50, 74% males, mean age ± SD: 2.71 ± 1.42 years, Data S1), randomly chosen from the sample of our earlier study (*18*). The dogs have been bred for and kept as experimental dogs by two research institutes, independently from the present study. The dogs from Institute 1 (N = 33) were kept in intra-specific groups of two to ten dogs and housed in indoor kennels with permanent access to outdoor runs. Dogs from Institute 2 (N = 17) were kept in same-sex pairs and housed in indoor kennels. Compared to the complex and stimulating environment of pet dogs living in human households, the kenneled dogs in our study experienced limited exposure to the outside world. The kennels at either facility had no environmental enrichment and interactions with humans were restricted to essential daily care, including feeding and cleaning, as well as a routine veterinary check-up once a month. In our previous study (*18*), we found no genetic or behavioral differences between the dogs from the two Institutes; thus, here, we analysed them together.

#### Behavioral phenotype

The dogs were tested using a standardised behavioral test; the detailed protocol for the test can be found in (*18*). For summary figure and table see Fig. S6 and Table S4.

The test battery took approximately 30 min to complete. Since the kenneled dogs did not have primary caretakers, their behavioral tests involved two experimenters. Experimenter 1, who was unfamiliar with the dog, conducted the test, while Experimenter 2 acted as the “owner", following a brief familiarization period, including gently talking to and petting the dog, taking the dog for a short walk, and offering food to the dog.

**Spontaneous Activity:** The owner (O) stands still while holding the dog on a 1.5–2 m leash, without engaging with it. The dog is free to explore within the leash's reach. No corrections or rewards are given. Experimenter 1 (E) remains at least 3 m away without interacting with the dog. The test lasts for 1 min.

**Greeting:** O stands still while holding the dog's leash. E approaches in a friendly manner, stopping just outside the dog's reach and waiting for 3 s. If the dog shows no aggression or avoidance, E steps forward and pets the dog’s head and back, then steps away for another 3 s before petting the dog again. If the dog avoids E without aggression, E remains outside the leash’s reach, crouches, and attempts to call the dog. If the dog approaches, E proceeds with petting as described. If the dog does not respond within 30 s, the test ends. In the case of aggression (e.g., growling, barking), E stays out of reach and speaks calmly to the dog for 10 s before terminating the test.

**Pendulum Test:** O stands behind the dog holding the leash and does not intervene.

Phase 1: To increase motivation, E moves a small piece of sausage in front of the dog three times before placing it on the ground for the dog to eat. The procedure is repeated in the opposite direction.

Phase 2: E swings a 10–12 cm sausage attached to a 30 cm string in front of the dog's nose, keeping it just out of reach for 30 s, before giving the dog a piece of sausage.

**Separation:** The dog is tethered on a ~3 m leash while O leaves and hides behind an object. After 1 min, E approaches and greets the dog (as in the Greeting test), then initiates play with a tug toy for 30 s before stepping back. After another minute, O returns, greets the dog, and also engages in play for 30 s.

**Ball Play:** The dog, either on a long (~3 m) leash or unleashed (for some family dogs), is encouraged by O to retrieve a tennis ball thrown a few meters away, repeated three times.

**Problem Solving:** The dog is presented with a small (22 × 14 × 15 cm) plastic cage with a narrow hole at the bottom, preventing direct access to a food reward inside. As pre-training, E places a piece of food outside the cage near the hole for the dog to eat. During the trial, O stands 1 m behind the dog, holding the leash and providing only verbal encouragement. E places a piece of food inside the cage and steps back. The dog has 60 s to retrieve the food by rolling or pushing the cage. The trial ends when the food is obtained or after 1 min (after which E gives the food to the dog). The test is repeated once.

**Bone Take Away :** O gives the dog a ham bone attached to a string and encourages chewing before stepping back. If the dog starts chewing, E approaches with an artificial hand (a plastic tube covered with a coat sleeve and a plaster-filled glove) and attempts to take the bone away in four escalating steps:

1. Petting the dog’s head and back three times.

2. Saying "please" while reaching for the bone.

3. Touching the bone for 3 s.

4. Pulling the bone away using the string while maintaining contact with the artificial hand.

The test is terminated if the dog is unmotivated to chew, allows E to take the bone, or displays severe aggression (e.g., snapping). If E cannot take the bone, O attempts it. The test is repeated once.

**Threatening Approach:** O holds the dog’s leash while E, standing 4–5 m away, approaches slowly with a slightly bent posture, maintaining direct eye contact and remaining silent. If the dog looks away, E attempts to regain attention with a cough or stomp. The test ends when E reaches the dog, the dog approaches E submissively/friendly, the dog persistently barks/growls, or the dog hides behind O. E then returns to the starting position, crouches, and calls the dog in a friendly manner.

**Umbrella:** O holds the dog’s leash while E approaches with a closed umbrella pointed at the dog. At ~1 m distance, E opens the umbrella, lifts it above her head, places it on the ground, and steps away. O then walks the dog past the umbrella. If the dog avoids it, O touches the umbrella and encourages the dog to approach.

**Lying on the Side:** O instructs the dog to lie down or gently places it in position, then turns the dog onto its side. If the dog resists, the test ends after 60 s. Otherwise, O holds the dog in position for 30 s, allowing petting and talking. If the dog rises before 30 s, the test restarts once. If the dog rises again, the test is terminated.

**Food Choice:** O holds the dog on a leash while E presents two plates—one containing a single piece of food and another with eight pieces—placing them 1.5 m apart. The dog is released to choose.

Phase 1: The dog chooses freely.

Phase 2: Before stepping back, E picks up the single piece, pretends to eat it with an audible "Hmm-mmm," then places it back on the plate before allowing the dog to choose.

**Hiding:** E holds the dog while O hides behind a large object (~15–20 m away). After 30 s, E releases the leash and says “Go!”. If the dog does not move within 5 s, E gently nudges it. If the dog does not seek O, O calls the dog. If the dog still does not approach, O returns while continuously calling.

**Spontaneous Activity 2:** The procedure is identical to the first Spontaneous Activity test.

*Coded behavioral variables*

All tests were video-recorded, and two behavioral variables were obtained. Inter-observer reliability was assessed in our previous study (*18*).

1. *Activity (%)*: The percent of the time the dogs spent with physical activity during the whole test. Activity was defined as moving the legs (locomotion) or active exploration (sniffing the ground or the objects) but did not include escape behaviors (i.e., actively retreating from the stimulus, struggling, and/or yanking the leash).
2. *Responsiveness* *(0-3)*: We reused the responsiveness scale scores introduced and published in (*18*). The score is calculated as the mean of 50 behavioral variables scored on a 0–3 scale in each subtest, where 0: no reaction, 3: intensive (positive) reaction. Responsiveness describes if and how far the individual was inclined to positively respond to different types of stimuli, including humans, objects, or the environment.

While high responsiveness means a more positive reaction (approach, interest, and attention) to the stimuli presented, low responsiveness could mean either indifference towards the stimuli or defence behaviors (i.e., immobility or active flight). With the activity variable, it was possible to differentiate between freezing (total immobility: low responsiveness, low activity) and flight (low responsiveness, high activity).

*Statistical analysis*

A hierarchical cluster analysis (Ward's method) based on responsiveness and activity was used to identify individuals with markedly different behaviors to determine the behavioral clusters (Fig. S1). For genotyping, prior to the GWAS analysis, the genotype data was filtered with the PLINK software based on SNP call rate (minimum 90%), minor allele frequency (minimum 1%), and based on Hardy-Weinberg equilibrium (α=0.05, with Bonferroni-correction applied for multiple testing). SNPs in complete LD were also removed. The GEMMA (Zhou & Stephens, 2012) software was used to implement the GWAS. For the GWAS analysis, a centered relatedness matrix was calculated with GEMMA using the default parameters except for the "-hwe" parameter, for which the default p-value (default: 0, i.e., no testing) was changed to 0.05 and for the missingness threshold, where the default (0.05) was changed to 0.1 in accordance with the genotype filtering step described above. In GEMMA, a univariate linear mixed model was fitted, and the Wald test was used to assess significance (α=0.05, with Bonferroni correction applied for multiple testing). In addition to the SNP effect, only an intercept was included in the model, but we also tested sex as a covariate.

##### Genotyping

Buccal swab samples were collected from the dogs in a non-invasive way by rubbing a pair of cotton swabs on the inner side of the dog's mouth (*17*). DNA was extracted from buccal swabs using the ethanol precipitation technique. The concentration of DNA solutions was determined using a NanoDrop 2000 spectrophotometer. Extracted DNA samples were stored at -20 C° long term after quantitation. Genotyping was done using the Illumina 170K Canine HD SNP array (Illumina, San Diego, CA-USA), which included 169,136 SNPs. Data can be found at <https://doi.org/10.57760/sciencedb.23921>.

##### Genome-wide association study

Call rates per individual ranged from 39,406 to 169,010; the average call rate was 89.1% (Data S1). Twelve animals had a call rate lower than 90%, which is most likely due to the inferior quality of the collected DNA sample; three of these samples had an extremely low call rate (23-27%), and these were not retained for further analysis. After SNP quality control, 82,409 SNPs were retained for GWAS. We carried out two GWAS analyses. First, we compared only the two extreme phenotype subclusters, Cluster 1 vs 4. Next, we compared the two large clusters (Cluster 1+2 vs Cluster 3+4).

**GWAS 1. Comparing Cluster 1 (freezing) and Cluster 4 (active-responsive)**

Assessment of population stratification and quality control: The estimated genomic inflation factor was close to one (1.01), measured across the whole genome. Therefore, we could conclude that no population stratification (other than that accounted for by the relatedness matrix included in the analysis) or systematic technical bias influenced our results. Fig. S7 shows the Q-Q plot between the expected and observed p-values after the - log_10_ transformation. This plot does not infer significant population stratification either.

Within a symmetric 200 Kb window surrounding the significant SNP, we calculated the fold-change difference between its p-value and the average p-value of nearby SNPs, finding that the lowest p-value was approximately 5 million times smaller.

The identified SNP – a transversion-type mutation (C/G) – can be found in the dbSNP database with the rs22253798 ID. It fell within the gene body of a known gene, namely the *potassium voltage-gated channel subfamily Q member 3* gene (*KCNQ3*; Ensembl ID: ENSCAFG00000001105). The SNP is located at intron 5 of the *KCNQ3-202* (ENSCAFT00000001691) and at the same intron of the *KCNQ3-203* (ENSCAFT00000086890) transcripts of the gene, but it did not overlap with the *KCNQ3-201* transcript (ENSCAFT00000064602). However, neither the SNP resolution nor the statistical power of this study is sufficiently high to conclude that any of the transcripts is of more interest than the other, and neither can the *KCNQ3*-201 transcript be ruled out completely. Therefore, the results are not discussed further at the transcript level.

Minor allele frequency of the significant SNP was 15.9% in the studied population. The linkage disequilibrium between the significant SNP and those in its vicinity was rather weak (r^2^_average_=0.355; r^2^_max_=0.667). The SNP's p-value showed a remarkable decrease (5,014,000x decrease) when compared to the p-values of the other SNPs in its vicinity.

Prior to the genome-wide association study, SNPs in complete LD were removed from the data. Following the GWAS analysis, we checked if the significant SNP had other SNP(s) in complete LD with it anywhere on the genome to avoid false interpretation of the results, but no such SNP was found.

**GWAS 2. Comparing Cluster 1+2 with Cluster 3+4**

After applying the same quality filters as for the first dataset, 65,035 SNPs were retained for the GWAS analysis. The results were evaluated and interpreted identically to GWAS 1. This analysis did not result in a PVE estimate that was significantly different from 0 (PVE=0.25 SE_PVE_=0.45), and the estimated genomic inflation factor deviated considerably from 1 as well ($\lambda=1.15$). The Q-Q plot indicated no population stratification. Furthermore, neither significant nor suggestive SNPs (i.e. SNPs close to the significance level) were identified in this study.

### *KCNQ3* expression and sequence in beagle dogs

##### i. *KCNQ3* expression analysis

**Samples:** Frontal cortex samples from five dogs were involved in the study (Table S1). Samples were obtained within four hours post-mortem from dogs that had been euthanised independently from this study and were voluntarily donated by the owners for research purposes. All the dissections and samplings were performed at ELTE by a veterinary anatomist. 100 mg fragments taken from the frontal cortex and brain stem regions were immediately immersed in RNAlater (Thermo Fisher) and were stored at  − 80°C following an overnight incubation at 4°.

**Protocol:** RNA isolation: Total cellular RNA was isolated from the RNAlater stabilized frontal cortex samples using TRIzol (Thermo Fisher) and following the manufacturer's protocol. Before immersing the tissue pieces in TRIzol, each piece was rinsed in 1 ml of sterile PBS in a new tube and centrifuged for 5 min on 500 g. TRIzol was added to the samples after the removal of the PBS. Homogenisation of the tissue pieces was carried out by an Ultra-Turrax homogeniser (Ika) under a fume hood to prevent inhalation of TRIzol. After homogenisation, the isolation process took place following standard procedures of TRIzol-based RNA isolation.

cDNA synthesis: 1 μg of each total RNA sample was reverse transcribed into cDNA, using the Maxima RevertAid cDNA Synthesis Kit (Thermo Fisher). Protocol was performed according to the manufacturer, using random hexamer primers. Prior to downstream applications, the cDNA samples were diluted tenfold in nuclease-free water and kept at -20°C, or for long-term storage, at -80°C.

qRT-PCR: Quantitative real-time PCR was performed on Roche LightCycle96 instrument, using Maxima SYBR Green qPCR Master Mix (2X) (Thermo Scientific). Two separate qRT-PCR reactions were executed with different normalisation controls (i.e. *ACTB*, *TUBA4A*) and all samples were run in quadruplicates in 96-well plates. Cycling parameters were as the following: 95°C x 10 min, then 40 cycles of 95°C x 15 sec, 60°C x 30 sec, and 72°C x 30 sec. The sample volume in each well was 10 μL with 0.5 μM final concentration of forward and reverse primers (Table S5). Data were analysed with R-based qRAT.

**Results:** The relative quantity of *KCNQ3* did not differ between the dogs as a function of behavior.

##### ii. RNA Sanger sequencing

**Samples:** Same samples as in the *KCNQ3* expression analysis (Table S1).

**Protocol:** Standard qRT-PCR reactions were performed, using the primer set shown in Table S6. We amplified four overlapping clones of the *KCNQ3* CDS. PCR fragments were cloned into the pGEM-T Easy Vector System (Promega, A1360) and transformed into *Escherichia coli* DH5α competent cells. After blue-white selection, 6-10 positive clones per PCR fragment were cultured overnight in a liquid medium. Following miniprep purification of the plasmid DNA, Sanger sequencing was performed using standard SP6 primers (Eurofins Gmbh).

**Results:** see Main text.

##### iii. Total RNA sequencing

**Samples:** Brain stem samples from four dogs, originally sequenced for (*29*) (Table S3). Data availability: <https://doi.org/10.57760/sciencedb.12821>.

**Protocol:** Sample preparation, sequencing, sequence data quality control and alignment to the canine reference genome (CanFam v3.1 reference genome version was used for compatibility reasons with the GWAS) was done as described in (*29*). Raw read coverage of the *KCNQ3* gene was calculated with the Rsubread R package (version 2.12.3), and the aligned data was visually examined using a locally installed version of the Integrative Genomics Viewer software. Raw sequence data is available at the *Sequence Read Archive*, under Bioproject ID: PRJNA939639.

**Results**: No differences were found in the *KCNQ3* gene as the function of the behavior (Fig. S4).

##### iv. DNA Sanger sequencing

**Samples:** Genomic DNA was isolated from buccal swabs of nine individuals, the same individuals as in the KCNQ3 expression analysis (Table S1) and four additional "freezing" animals (Table S7).

**Protocol:** DNA isolation: Genomic DNA was isolated from buccal swabs using the following procedure. Buccal swabs were placed in labelled 1.5 mL microcentrifuge tubes. Each sample was lysed with 630 µL of TNES buffer (10 mM Tris, pH 7.5; 400 mM NaCl; 100 mM EDTA; 0.6% SDS) and 6 µL of Proteinase K (20 mg/mL), followed by gentle inversion to mix. The samples were incubated at 55°C overnight with occasional gentle inversion. After incubation, 200 µL of 5 M NaCl was added, and the samples were shaken vigorously for 20 seconds before centrifugation at 12,000-14,000 rpm for 5-10 minutes at room temperature. The supernatant (750 µL) was carefully transferred to new tubes, avoiding the pellet. An equal volume of ice-cold 100% ethanol (750 µL) was added, and the mixture was gently inverted to precipitate the DNA. The samples were centrifuged at 5000 x g for 20-25 minutes at 4°C. The supernatant was removed, and the DNA pellet was washed with 500-1000 µL of cold 100% ethanol, followed by centrifugation at 5000 x g for 5 minutes at 4°C. The ethanol was removed, and the pellet was washed with 70% ethanol in the same manner. The DNA pellets were air-dried for 10-30 minutes and resuspended in 100-200 µL of TE buffer (10 mM Tris-HCl, pH 8.0; 0.1 mM EDTA). For higher purity, resuspended DNA was treated with RNase A (100 µg/mL) and phenol:chloroform:isoamyl alcohol (25:24:1) (PanReac - A2279), followed by re-precipitation with sodium acetate and ethanol.

Genotyping PCR: For genotyping, we used genomic DNA from all individuals and amplified a 232 bp long portion of the *KCNQ3* exon 1 with Thermo Scientific Platinum SuperFi II DNA polymerase (Catalog number 12369010, protocol according to manufacturer), using forward KCNQ3-Cf-F1 (5'- ACGAGGAGCGCAAAGTGG-3') and reverse KCNQ3-GT-R (5'- AGGGCGTCGTAGATCAAAGTTTGG-3') primers. The amplified fragments were then sequenced via Sanger sequencing.

**Results:** Sequence data were not different across dogs.

### 3. Validating the *KCNQ3*'s role in stress regulation in a fish model

##### Mutagenesis

Alt-R crRNAs targeting exons 2, 3 and 5 of *kcnq3* were ordered from Integrated DNA Technologies (IDT, Table S8). After annealing them with the Alt-R tracrRNAs (catalog no. 1072532, IDT) and the incubation of the crRNA:tracrRNA complexes with Cas9 protein, the resulting RNPs were injected into one-cell-stage embryos using glass capillaries. Final concentrations were 5.7 µM for each crRNA:tracrRNA hybrid and 3.47 µM for Cas9 protein.

##### Phenotyping

***Tactile startle (TS) paradigm***

TS experiments were carried out on 6-well plates containing 3 ml of fish water, placed in a Zantiks MWP unit. The test consisted of a baseline and a stimulus period, both one-minute-long, in which the subjects received no or repeated tactile stimuli in standard 1-second intervals, respectively. Zebrafish either tend to startle, make escape manoeuvres or generally enhance or decrease their locomotion in response to tactile stimuli. Changes in these trends can be considered as differences in the fear and accompanied arousal state of the subjects (Colwill & Creton, 2011). Two-minute-long tests were video recorded with the MWP unit, and the total distance moved in 250 millisecond time bins was calculated using Ethovision.

***The swimming plus-maze (SPM) test***

The SPM test is conducted in a plus-shape platform that consists of two deeper and two shallower arms. The approach-avoidance behavior of zebrafish in the test is driven by the trade-off between explorative motivation and anxiety-like states in the proximity of the threatening water surface in the shallow arms (*33*). Ten-minute long tests were video recorded using the MWP unit, and the frequency and latency to enter and time spent in the shallow arms were measured. The total distance moved by the fish was also measured to represent general locomotion. Measures of shallow arm activity were summarised to an anxiety score using a formula of anxiety score = (scaled frequency + scaled time - scaled latency) * (-1), which positively represents the anxiety-like state of zebrafish.

##### Genotyping

Figure S8 presents the CRISPR-based F0 mutagenesis of zebrafish *kcnq3*. Target sequences were (with PAM sequence in bold): exon 2 – 5’-AGGGAAGTGGACTTCAAAAG**GGG**-3', exon 3 – 5’-ACGAGAAGGATTCGGCTCAC**TGG**-3', exon 5 – 5’-TGATTACGTACGGCAACCAC**CGG**-3'. Primers used for the PCR amplification of genomic fragments can be found in Table S9.

##### Statistical analysis

Statistical analysis was done using the R statistical environment (R Core Team, 2021). To test the relationship between the genotype, the tactile stimuli and the locomotion of animals, we conducted two-way repeated-measure ANOVA models, using the genotype and time bin as fixed independent variables and the ID of animals as repeated factors. Main effect models were done using the *lme4* package, and post-hoc comparisons were made using the *emmeans* package. To test the relationship between the genotype and anxiety-like states and locomotion, we conducted student *t*-tests.

**Fig. S1.**

**Behavioral categorization of dogs.** **(a)** Cluster analysis grouped the dogs into two major clusters which were further divided into two-two sub-clusters. The four sub-clusters showed significant differences in **(b)** total activity and **(c)** responsiveness scores.

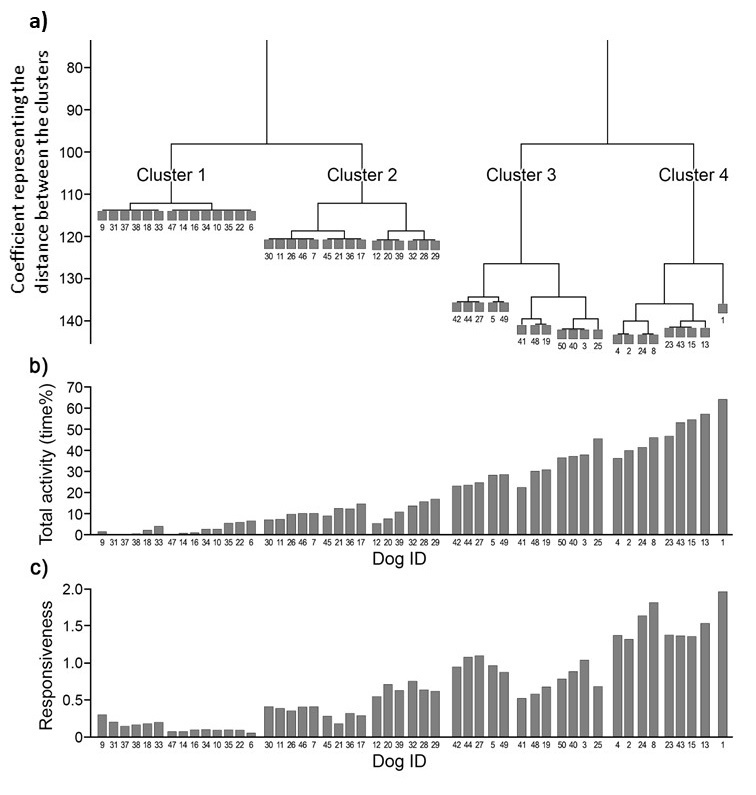

**Fig. S2.**

**Manhattan plot of the genome-wide association analysis of the freezing behavior in beagle dogs.** X-axis: chromosomes and genomic positions; y-axis: $-{log}_{10}$ transformed p-values. The red line represents the Bonferroni-corrected alpha value (α=0.05; α_corrected_=6.2). The red dots represent the significant SNP and those SNPs that fall within its proximity.

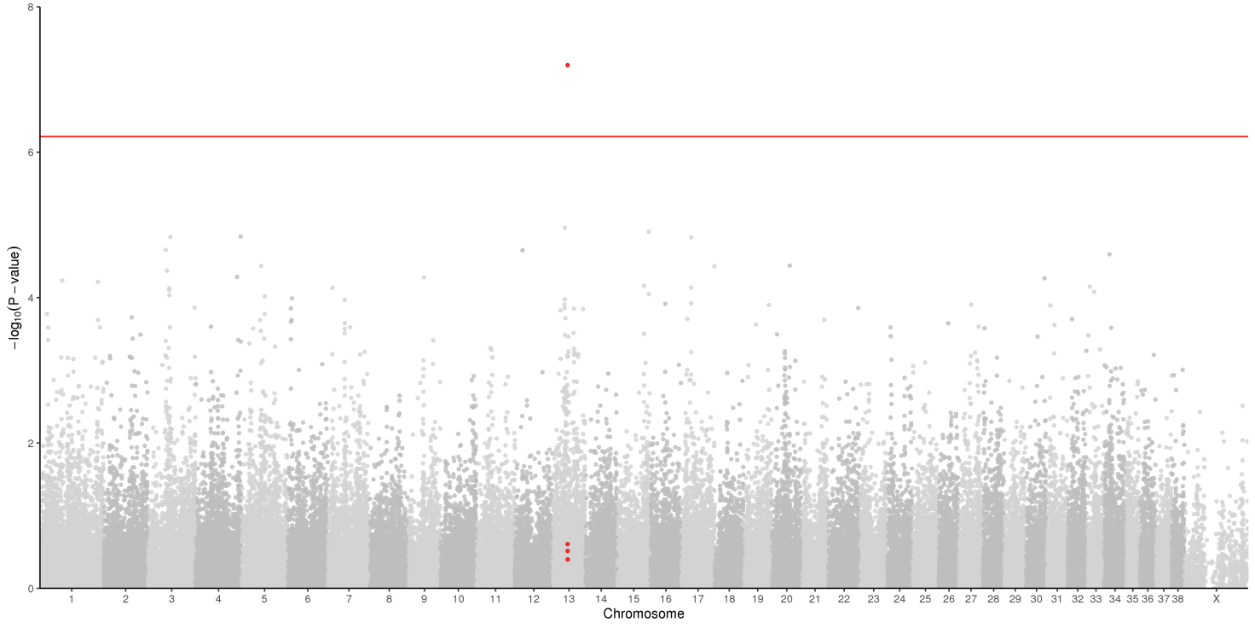

**Fig. S3.**

**qRT-PCR** results using (A) *ACTB* and (B) *TUBA4A* as the normalisation control. Only the last number of the brain ID-s is shown in the figures (see Table S2); letters indicate several samples from the same brain. In Fig. S5A, sample 1b was excluded because the cDNA sample did not amplify the *KCNQ3*.

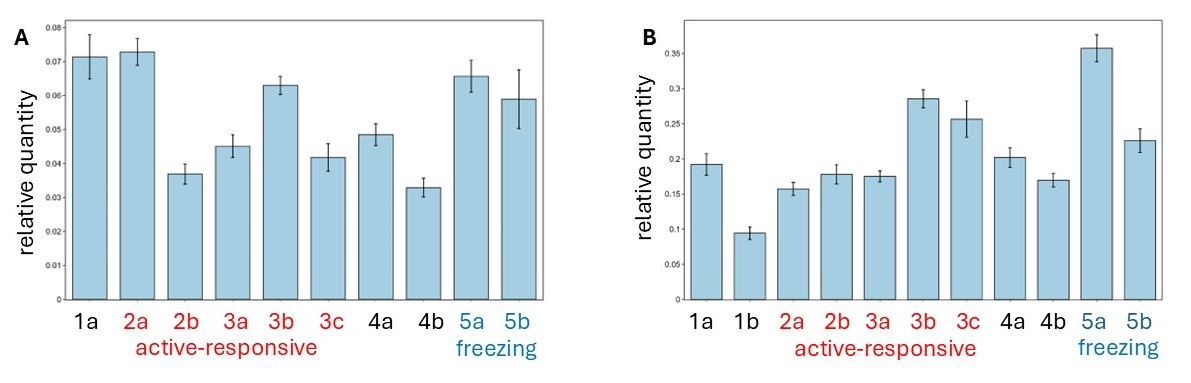

**Fig. S4.**

Coverage tracks of the *KCNQ3* gene (a), the 3' end of the gene is enlarged (b).

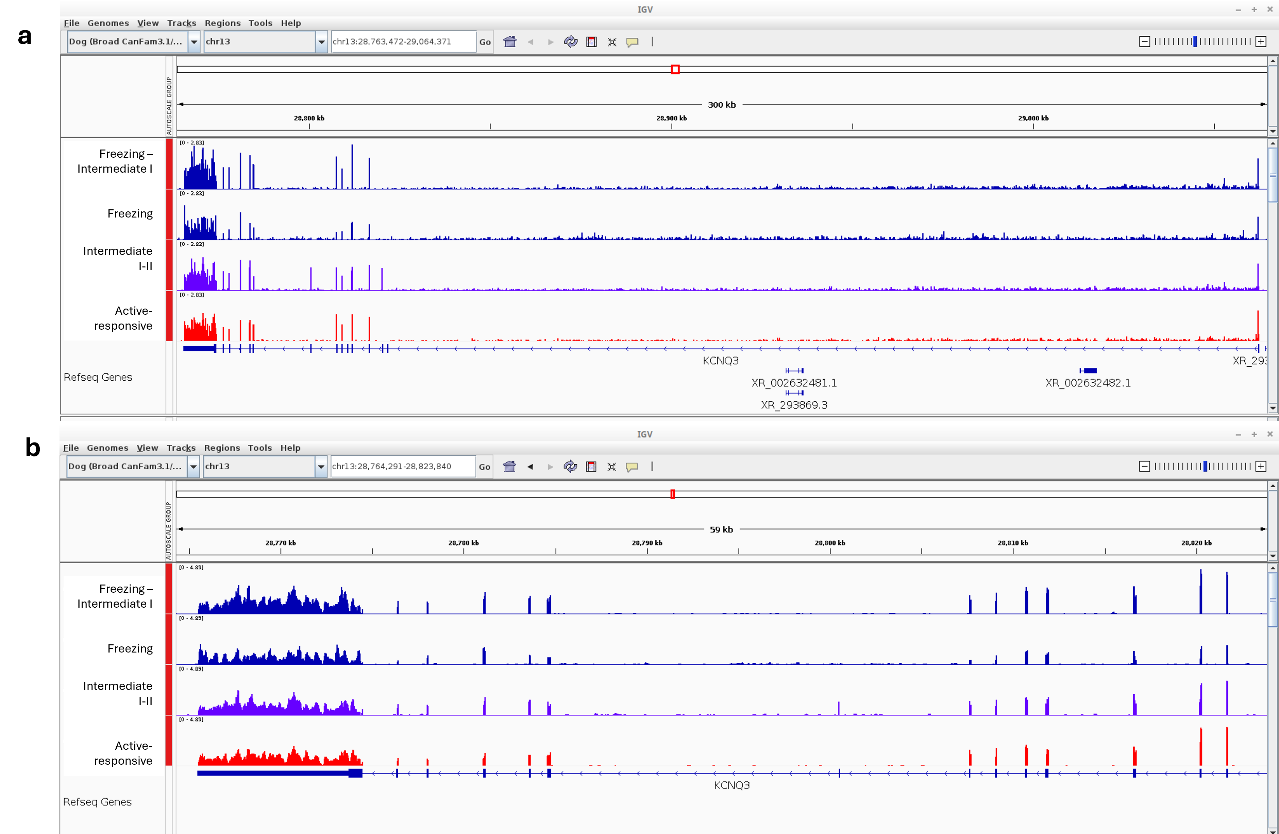

**Fig. S5**.

Similarity of dog and zebrafish *KCNQ3* orthologs**.** Dot plot of a BLAST protein sequence comparison between the KCNQ3 proteins of *Canis familiaris* (transcript: A0A8I3N9A5) and *Danio rerio* (A0A8M9PQX0).

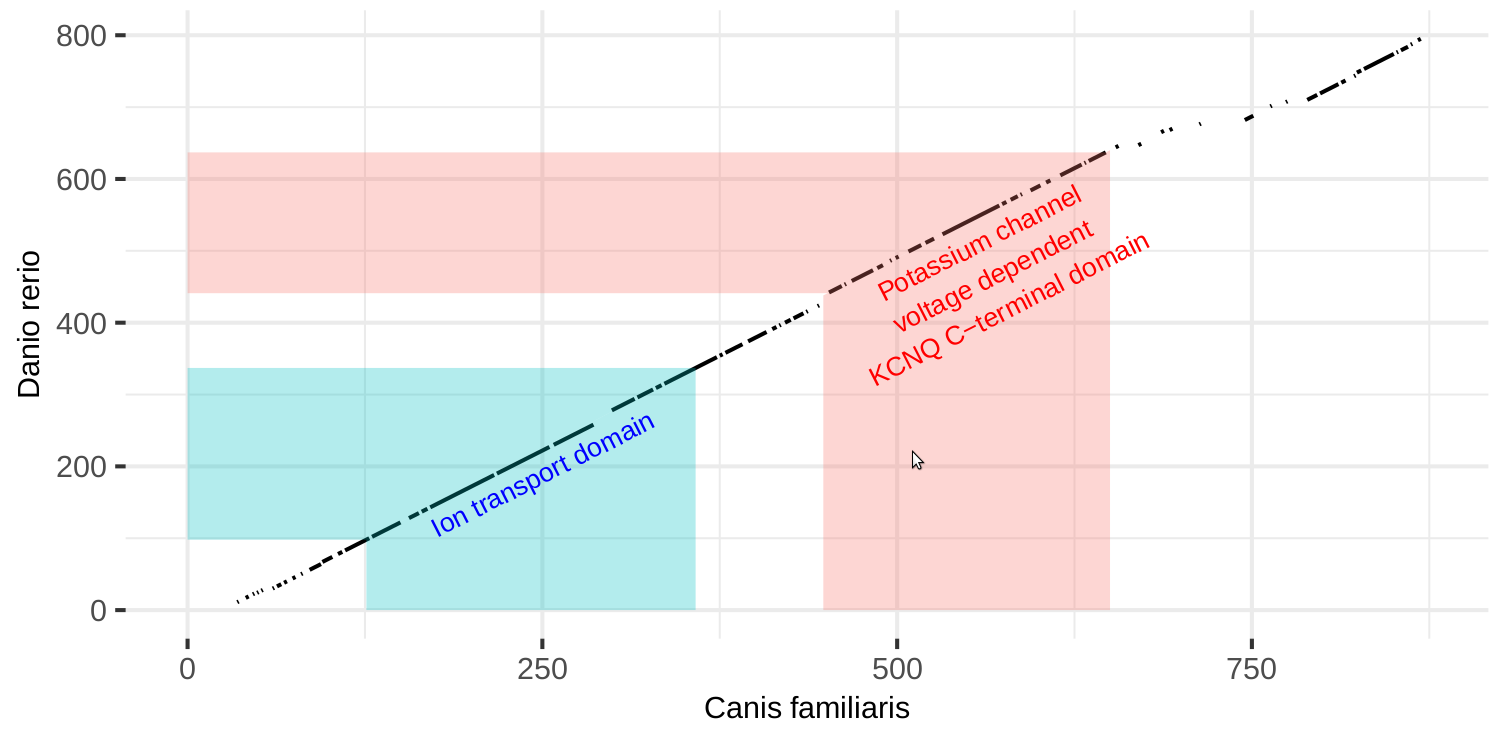

Fig. S6.

Subtests of the personality test applied in the study (*18*). (The persons presented in the images are coauthors EK and BT. They both provided written consent to publication.)

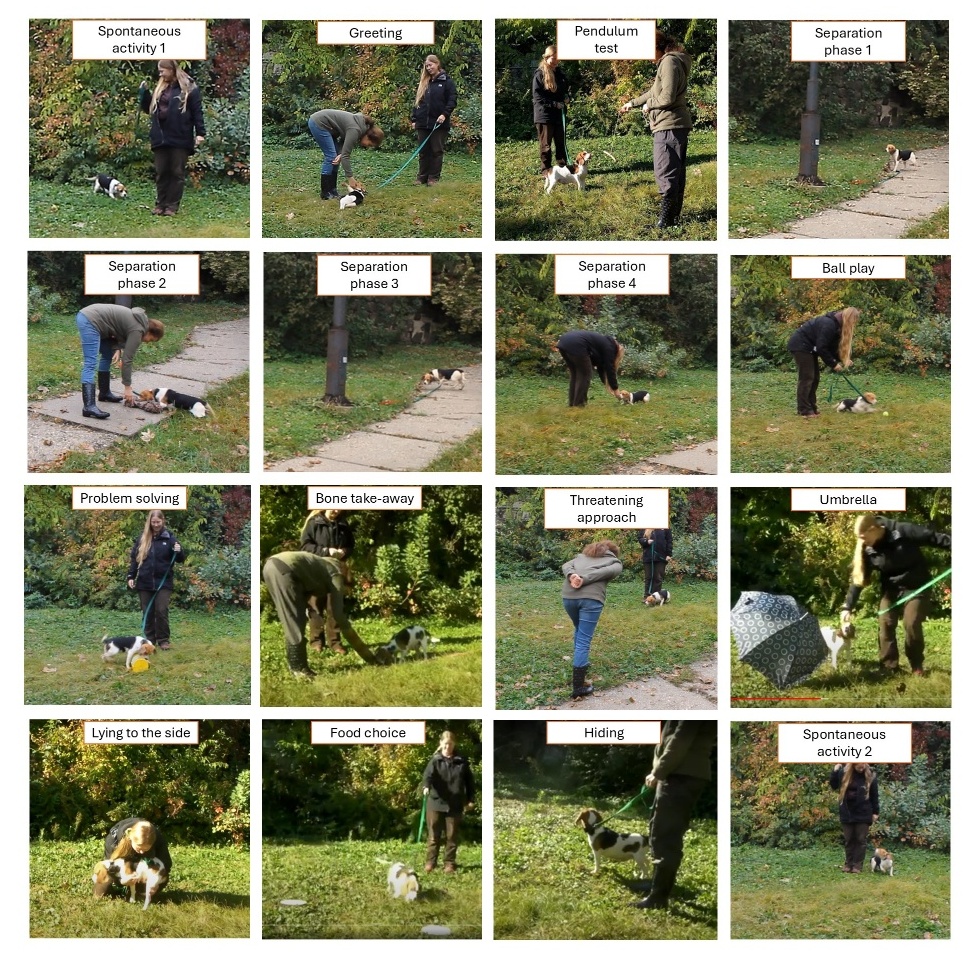

**Fig. S7.**

Q-Q plot of expected vs. observed GWAS p-values. P-values are shown after –log_10_ correction).

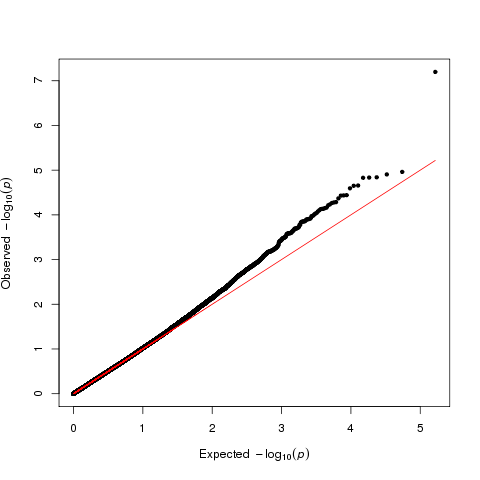

**Fig. S8.**

CRISPR-based F0 mutagenesis of zebrafish *KCNQ3*. **(a)** Schematic representation of the zebrafish *kcnq3* gene, denoting the binding position of the crRNAs injected. **(b-d)** Sequenograms show the efficiency of CRISPR-based Cas9 targeting in exons 2 **(b)**, 3 **(c)** and 5 **(d).** The binding position and orientation of the respective crRNAs within the exons are indicated. The result of Sanger sequencing suggests the presence of multiple indels upon the injection of the Cas9 RNPs.

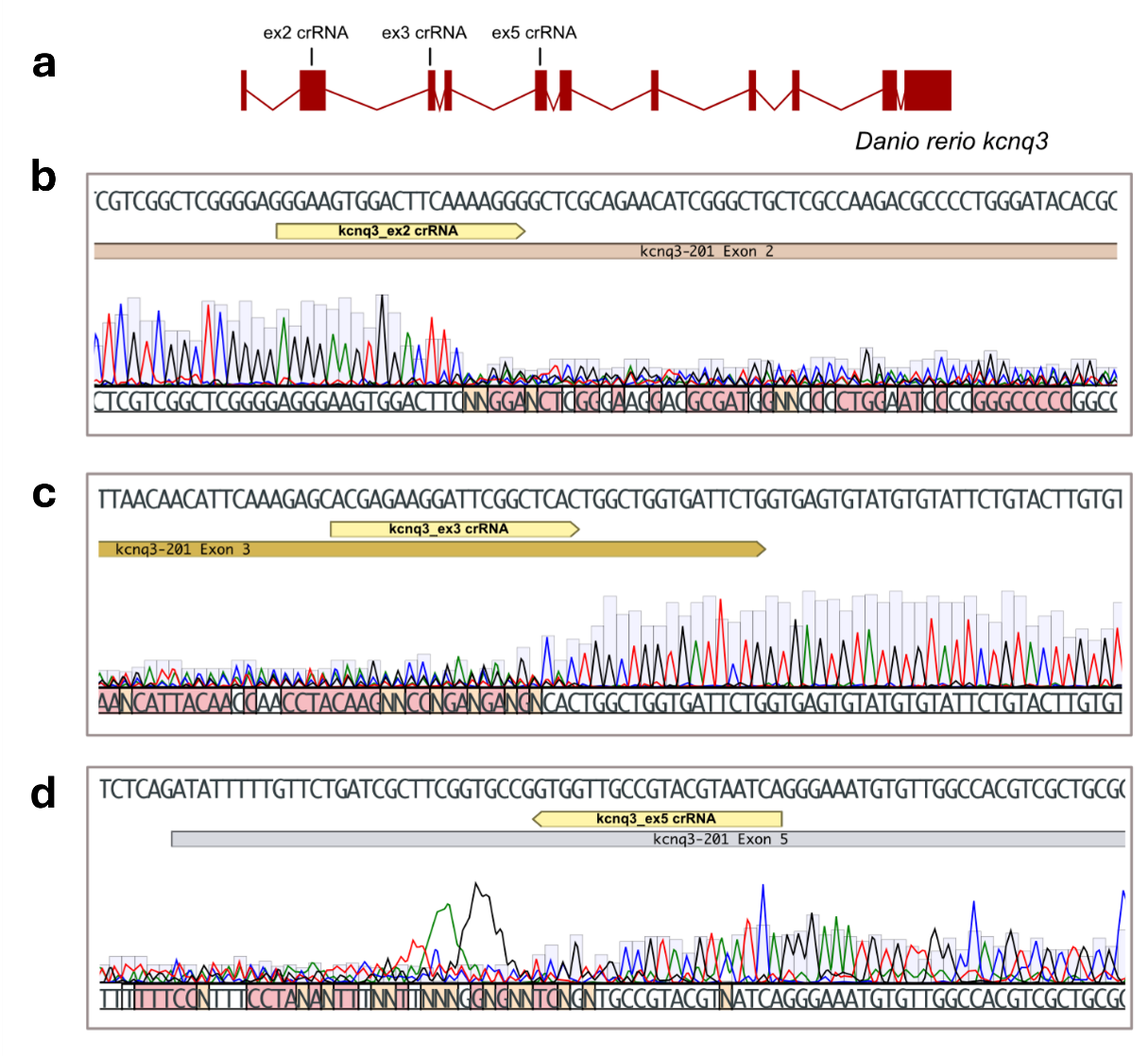

**Table S1.**

The behavior of dogs in the *KCNQ3* expression analysis and the RNA Sanger sequencing. *These dogs were not included in the cluster analysis, but were included in (*18*), therefore, their behavior was known.

| Cluster ID* | Cluster name | Brain ID | DNA ID | activity (%) | responsiveness |
| --- | --- | --- | --- | --- | --- |
| 1 | Freezing | 180420-5 | KK3635 | 4.08 | 0.19 |
| 2-3 | Intermediate I-II. | 180420-1 | KK3637 | 20.30 | 0.21 |
| 3 | Intermediate II. | 180420-4 | KK3634 | 16.54 | 1.17 |
| 4 | Active-responsive | 180420-2 | KK3636 | 58.10 | 1.44 |
| 4 | Active-responsive | 180420-3 | KK3633 | 40.65 | 1.51 |

**Table S2.**

Point mutations identified in the *KCNQ3* coding sequence of the "freezing" dog (brain ID: 180420-5). *indicates missense mutations.

| **Amino acid change** | **Reference codon** | **New codon** |
| --- | --- | --- |
| T685 | ACC | ACT |
| K732N* | AAG | AAT |
| C773 | TGT | TGC |
| S774 | TCA | TCG |

**Table S3.**

The behavior and CPM (counts of reads per gene per million sequenced reads) of dogs in total RNA sequencing. *These dogs were not included in the cluster analysis but were included in (*18*), therefore, their behavior was known.

| Cluster ID* | Cluster name | Brain ID | DNA ID | Tandon et al. (2024) (*29*) ID | activity (%) | responsiveness | CPM |
| --- | --- | --- | --- | --- | --- | --- | --- |
| 1 | Freezing | 180420-5 | KK3635 | SRX19523800 | 4.08 | 0.19 | 22.2 |
| 1-2 | Freezing - Intermediate I. | 171102-2 | KK3421 | SRX19523791 | 6.95 | 0.54 | 50.6 |
| 2-3 | Intermediate I-II. | 180420-1 | KK3637 | SRX19523799 | 20.30 | 0.21 | 33.0 |
| 4 | Active-responsive | 171102-1 | KK3416 | SRX19523790 | 52.02 | 1.63 | 44.1 |

**Table S4.**

Summary of the test protocol. O: owner, E: experimenter. The test was preceded by a short (approx. 5 min.) familiarisation with one of the experimenters, who then played the part of the owner. The location of the test was set to match the dogs' housing conditions (outdoors for dogs from Institute 1 and indoors for dogs from Institute 2). See also Fig. S6.

| **Subtest** | **Protocol** |
| --- | --- |
| Spontaneous activity | The dog (on a 1.5-2 m leash) is free to explore its environment (1 min) |
| Greeting | E approaches the dog in a friendly manner and tries to pet it |
| Pendulum test | E swings a sausage in front of the dog's nose just out of its reach |
| Separation - Phase 1 | The dog is left alone for 1 min |
| Separation - Phase 2 | E approaches the dog and tries to play with it |
| Separation - Phase 3 | The dog is left alone for 1 min |
| Separation - Phase 4 | O comes back and tries to play with the dog |
| Ball play | O plays with the dog using a tennis ball |
| Problem-solving | The dog has 1 min to obtain a piece of food from a small cage (2 trials) |
| Bone take away | The dog is given a large bone, which the E tries to take away (2 trials) |
| Threatening approach | E approaches the dog slowly, in a threatening manner |
| Umbrella | E opens an umbrella close (~ 1 m) to the dog |
| Lying on the side | O tries to lay the dog to its side and keep it in this position (1 min) |
| Food choice | The dog can choose between 1 and 8 pieces of food, the former was preferred by the E |
| Hiding | O leaves the dog with E for 30 sec then the dog is allowed to go free |
| Spontaneous activity 2 | Same as Spontaneous activity |

**Table S5.**

Primers used to amplify *KCNQ3* cDNA sequences from dog brain samples.

| **Primer name** | **Primer sequence (5’-3')** |
| --- | --- |
| KCNQ3-Cf-F1 | ACGAGGAGCGCAAAGTGG |
| KCNQ3-Cf-R1 | ggagacgccgattaaggaga |
| KCNQ3-Cf-F2 | atgtgccagaggttgatgca |
| KCNQ3-Cf-R2 | cgatgtggatgctctggcta |
| KCNQ3-Cf-F3 | gcaggacatctcgacatgct |
| KCNQ3-Cf-R3 | ctccctcatccagcttgacc |
| KCNQ3-Cf-F4 | Tttttgcccacgacccagt |
| KCNQ3-Cf-R4 | cgtcccggtccaatcatgat |

**Table S6.**

Primers used for qRT-PCR.

| **Primer name** | **Primer sequence (5’-3')** |
| --- | --- |
| KCNQ3-Cf-qPCR-F | TCAGGCTGCCTGGAGGTATTATGCTACC |
| KCNQ3-Cf-qPCR-R | GCTACCACGAGGATTAGAAAGGCGAACC |
| ACTB-Cf-qPCR-F | CGGCATCCATGAAACTACCTTCAACTCC |
| ACTB-Cf-qPCR-R | GCGATGATCTTGATCTTCATTGTGCTGG |
| TUBA4A-Cf-qPCR-F | GGGAAGGAGGATGCTGCTAACAA |
| TUBA4A-Cf-qPCR-R | TTCTTGCCGTAGTCAACCGAGAG |

**Table S7.**

The behavior of four "freezing" dogs (Cluster 1), included in DNA Sanger sequencing in addition to dogs in Table S3.

| DNA ID | Cluster | Dog ID in Fig. S2 | activity (%) | responsiveness |
| --- | --- | --- | --- | --- |
| KK2977 | 1 | 31 | 0 | 0.20 |
| KK3117 | 1 | 37 | 0 | 0.14 |
| KK3118 | 1 | 38 | 0.15 | 0.16 |
| KK3128 | 1 | 47 | 0 | 0.71 |

**Table S8.**

Guide RNAs (gRNAs) targeting exons 2, 3, and 5 of the kcnq3 gene.

| **gRNA name** | **gRNA sequence** |
| --- | --- |
| kcnq3_ex2-gRNA | AGGGAAGTGGACTTCAAAAG |
| kcnq3_ex3-gRNA | ACGAGAAGGATTCGGCTCAC |
| kcnq3_ex5-gRNA | TGATTACGTACGGCAACCAC |

**Table S9.**

Primers used to amplify targeted regions of the zebrafish *kcnq3* locus.

| **Primer name** | **Primer sequence (5’-3')** |
| --- | --- |
| KCNQ3-Dr-ex2_F | GGGATCAGGTCCAGAAATGCCGC |
| KCNQ3-Dr-ex2_R | TGGTAGAGCAGCGCCCATCCTC |
| KCNQ3-Dr-ex3_F | TGTTTTGGGATGTCTGATTCTGTCG |
| KCNQ3-Dr-ex4_R | CGGTTTGCGGGCGAATTTGAGC |
| KCNQ3-Dr-in4_F | TGCCGTTTGTGCTTATTTTGTATG |
| KCNQ3-Dr-ex5_R | CTCCGCGTCTGTCCATTCGCAG |

**Movie S1.**

Video about a dog showing freezing behavior (Cluster 1). See also here: <https://youtu.be/i2zJ0C-5k8w>.

**Movie S2.**

Video about an active-responsive dog (Cluster 4). See also here: <https://youtu.be/shK3Nk4MAPU>.

Data S1.

The 50 genotyped dogs’ demographics, behavior and SNP call rates.

| **DNA ID** | **Age (year)** | **Sex** | **Moving %** | **Responsiveness** | **Cluster** | **Called SNP N** | **SNP %** |
| --- | --- | --- | --- | --- | --- | --- | --- |
| KK2977 | 4.2 | male | 0.0 | 0.2 | 1 | 168739 | 99.77 |
| KK3117 | 3.3 | male | 0.0 | 0.1 | 1 | 168553 | 99.66 |
| KK3128 | 1.5 | male | 0.0 | 0.1 | 1 | 168841 | 99.83 |
| KK3118 | 1 | female | 0.1 | 0.2 | 1 | 168799 | 99.8 |
| KK1232 | 2 | male | 0.5 | 0.1 | 1 | 168299 | 99.51 |
| KK1228 | 2 | male | 0.7 | 0.1 | 1 | 168959 | 99.9 |
| KK919 | 2 | male | 1.1 | 0.3 | 1 | 168978 | 99.91 |
| KK1221 | 2.4 | male | 1.9 | 0.2 | 1 | 166839 | 98.64 |
| KK3114 | 1.5 | male | 2.3 | 0.1 | 1 | 168865 | 99.84 |
| KK921 | 3 | male | 2.3 | 0.1 | 1 | 168930 | 99.88 |
| KK3055 | 1.2 | female | 3.7 | 0.2 | 1 | 168725 | 99.76 |
| KK3115 | 1.5 | male | 5.2 | 0.1 | 1 | 168803 | 99.8 |
| KK1231 | 2 | male | 5.6 | 0.1 | 1 | 168974 | 99.9 |
| KK911 | 3 | male | 6.4 | 0.1 | 1 | 168844 | 99.83 |
| KK923 | 2 | male | 5.1 | 0.5 | 2 | 168172 | 99.43 |
| KK2974 | 4.2 | male | 6.8 | 0.4 | 2 | 45224 | 26.74 |
| KK922 | 3 | male | 7.1 | 0.4 | 2 | 168804 | 99.8 |
| KK1225 | 2 | male | 7.4 | 0.7 | 2 | 39406 | 23.3 |
| KK3125 | 1.5 | female | 8.8 | 0.3 | 2 | 155390 | 91.87 |
| KK2976 | 1.2 | female | 9.5 | 0.4 | 2 | 162963 | 96.35 |
| KK3127 | 1.8 | male | 9.8 | 0.4 | 2 | 165218 | 97.68 |
| KK912 | 2 | male | 9.8 | 0.4 | 2 | 168415 | 99.57 |
| KK3119 | 1 | female | 10.5 | 0.6 | 2 | 92712 | 54.82 |
| KK3116 | 2.6 | male | 12.1 | 0.3 | 2 | 161084 | 95.24 |
| KK1236 | 3 | male | 12.2 | 0.2 | 2 | 138994 | 82.18 |
| KK3054 | 2.5 | male | 13.4 | 0.8 | 2 | 168963 | 99.9 |
| KK1235 | 2 | male | 14.4 | 0.3 | 2 | 168508 | 99.63 |
| KK2971 | 3.5 | female | 15.5 | 0.6 | 2 | 167455 | 99.01 |
| KK2973 | 3.5 | male | 16.6 | 0.6 | 2 | 159310 | 94.19 |
| KK3121 | 1.1 | male | 22.3 | 0.5 | 3 | 113556 | 67.14 |
| KK3122 | 4.1 | male | 22.9 | 0.9 | 3 | 156608 | 92.59 |
| KK3124 | 1.6 | female | 23.2 | 1.1 | 3 | 168970 | 99.9 |
| KK2978 | 2.5 | female | 24.5 | 1.1 | 3 | 149675 | 88.49 |
| KK910 | 2 | male | 28.0 | 1.0 | 3 | 168392 | 99.56 |
| KK3130 | 6.5 | female | 28.4 | 0.9 | 3 | 169010 | 99.93 |
| KK3129 | 1.6 | male | 29.8 | 0.6 | 3 | 148987 | 88.09 |
| KK1226 | 2 | male | 30.5 | 0.7 | 3 | 167419 | 98.98 |
| KK3131 | 6.6 | female | 36.3 | 0.8 | 3 | 107680 | 63.66 |
| KK3120 | 1.1 | male | 37.0 | 0.9 | 3 | 43510 | 25.72 |
| KK903 | 4 | male | 37.8 | 1.0 | 3 | 168864 | 99.84 |
| KK2975 | 3.4 | female | 45.4 | 0.7 | 3 | 111423 | 65.88 |
| KK904 | 6 | male | 36.1 | 1.4 | 4 | 166656 | 98.53 |
| KK902 | 6 | male | 39.7 | 1.3 | 4 | 168991 | 99.91 |
| KK2989 | 1.1 | female | 41.3 | 1.6 | 4 | 160267 | 94.76 |
| KK918 | 2 | male | 45.9 | 1.8 | 4 | 168800 | 99.8 |
| KK2988 | 4.8 | male | 46.6 | 1.4 | 4 | 168909 | 99.87 |
| KK3123 | 4.1 | male | 52.9 | 1.4 | 4 | 106518 | 62.98 |
| KK1227 | 2.8 | male | 54.3 | 1.4 | 4 | 168991 | 99.91 |
| KK1153 | 3 | female | 57.0 | 1.5 | 4 | 105452 | 62.35 |
| KK901 | 3 | male | 64.0 | 2.0 | 4 | 162288 | 95.95 |
